# Early HIV-1 maturation drives Env clustering and fusion competence

**DOI:** 10.64898/2025.12.11.693442

**Authors:** Irene Carlon-Andres, Victoria Garcia-Giner, David J. Williamson, Ravi Teja Ravi, Audrey Le Bas, Philip N. Ward, Tobias Starling, Lina El Hajji, Arnaud Gautier, Sabrina Simoncelli, Michael Grange, Maud Dumoux, Sergi Padilla-Parra

**Author notes:** Corresponding authors at;.

## Abstract

Human immunodeficiency virus type 1 (HIV-1) particles are initially released from the cell in an immature state that cannot cause infection. The proteolytic processing of the Gag structural polyprotein into its individual components triggers a dramatic structural reorganisation that produces fully mature and infectious virions. HIV-1 maturation remodels the Gag lattice to enable membrane fusion, but how early proteolytic steps regulate the envelope glycoprotein (Env) organisation is unknown. Although cryo-electron tomography has defined structural intermediates of maturation, it has not resolved how Env conformers relate to specific early maturation states. Here, we show that partial Gag cleavage, prior to capsid formation, allosterically reorganises the membrane-proximal lattice, promoting Env clustering and fusion-competent conformations. Using native virus preparations combined with cryogenic fluorescence-lifetime imaging correlated to cryo-electron tomography, we map Env conformational signatures onto defined Gag-processing intermediates. This reveals an unrecognised early maturation stage in which incomplete Gag processing primes Env for entry, shifting current models by positioning functional Env activation upstream of core maturation. These results establish that early lattice remodelling is a key determinant of HIV-1 fusion competence and expose a previously inaccessible, intervention-sensitive step with implications for therapeutic and vaccine strategies targeting maturation–entry coupling.

## Main

Human Immunodeficiency Virus type 1 (HIV-1) undergoes a tightly regulated maturation process essential for viral entry and productive infection. Nascent virions assemble at the plasma membrane through multimerization of the Gag and GagPol precursor polyproteins and incorporation of the envelope glycoprotein (Env), resulting in immature particles characterized by a Gag spherical lattice underneath the viral membrane^1^. Gag comprises the matrix (MA), capsid (CA), nucleocapsid (NC), and p6 domains, as well as short spacer peptides SP1 and SP2, whereas GagPol additionally includes the viral protease (PR), reverse transcriptase (RT), and integrase (IN)^1^. During particle budding, protease dimerization triggers sequential cleavage of the precursor proteins^2^. Initial cleavage separating SP1 from NC allows NC–RNA complex condensation. A subsequent cleavage at the MA–CA junction releases MA and detaches the CA–SP1 region from the membrane, followed by CA-SP1 processing^3^. Gag processing increases Env mobility inducing its redistribution at the viral membrane^4–7^ and drives formation of the conical capsid that encloses and protects the viral genome, resulting in major structural rearrangements that transform the immature particle into the mature virion. The ordered cleavage of Gag/Gag-Pol and the remodelling of the lattice into a mature capsid are essential for infectivity^8–10^. Protease inhibitors disrupt these steps and have therefore become a central component of current HIV therapy^11^. Despite major advances, we still lack a complete structural and mechanistic understanding of how sequential Gag processing coordinates Env redistribution and the acquisition of fusion and infectivity. Defining these transitions is crucial for the rational design of next-generation treatments.

Light microscopy methods have contributed to understanding HIV-1 maturation by providing dynamic, quantitative information about the timing and regulation of Gag processing, employing fluorescent proteins or tags fused to the accessory protein Vpr^12,13^ or to the structural polyprotein Gag^14–18^. However, they have not directly linked single-particle fluorescence readouts of molecular maturation states to virion structure and downstream fusion and infection efficiencies. We have developed a modular FRET–based biosensor enabling rapid, on-demand generation of multiple spectrally distinct FRET pairs, providing greater flexibility than fixed fluorescent protein pairs and facilitating multiplexing with additional markers^19^. Our biosensor is inserted between MA and CA subunits of Gag, providing a direct read-out of matrix release from the Gag lattice during maturation.

Vitrification by rapid-cooling preserves biological samples in a near-native, frozen-hydrated state, enabling structural analysis with cryo-electron tomography (cryoET)^20–23^. Similarly, cryo-fluorescence microscopy is an increasingly used method to correlate features of interest in vitrified samples (including in cells and tissues). However, current approaches lack functional information to guide structural analysis^24^. Fluorescence lifetime imaging microscopy (FLIM) measures the fluorescence decay times after fluorophore excitation, revealing details about its micro-environment, such as proximity to a quencher fluorophore, at the molecular level. FLIM detects FRET (Förster resonance energy transfer) between fluorophores separated up to ∼10 nm, relying solely on donor fluorescence and avoiding artifacts of intensity-based measurements^25–27^. This opens the possibility of performing correlative studies that directly link specific protein dynamic functions to their structural context.

We use FRET-FLIM to capture conformational changes during HIV-1 maturation under physiological and cryogenic conditions, enabling correlative analysis of transient states with molecular details using cryo-ET. Our optimized virus production preserves native, full-length JR-FL Env on particle surfaces, allowing direct links between HIV-1 core morphology, Env organization, and functional competence. We show that early protease-mediated matrix release from the Gag lattice triggers Env clustering, which precedes capsid formation and enhances fusion but not infectivity. Targeting early steps of Gag processing or Env clustering during maturation could provide new avenues for antiviral intervention. Beyond HIV-1, this approach establishes a general framework to couple protein activity dynamics with structural context in intact, near-native systems.

## Multi-spectral and lifetime FRET-FLIM biosensor of HIV-1 maturation

Newly assembled HIV-1 particles are released from host cells in an immature state and must undergo viral protease-mediated processing to become fully functional mature virions. To track the HIV-1 maturation process, we engineered a genetically encoded FRET-based biosensor incorporated into viral particles (Fig. 1a). The FRET pair is introduced between the matrix and the capsid domains of the precursor protein Gag, flanked by the SQNYPIVQ protease cleavage site naturally present between matrix and capsid^28^. On-demand FRET occurs between an intrinsically fluorescent protein (e.g. eGFP) serving as the donor and FAST, a scaffold protein, serving as the acceptor^29–31^. FRET is triggered on demand by addition of a synthetic fluorogen that permeates the viral membrane and binds specifically to FAST. Therefore, protease processing of the MA/CA interphase releases the FRET pair components, inducing a change in the distance which results in a decrease of FRET signal.

**Fig. 1.**
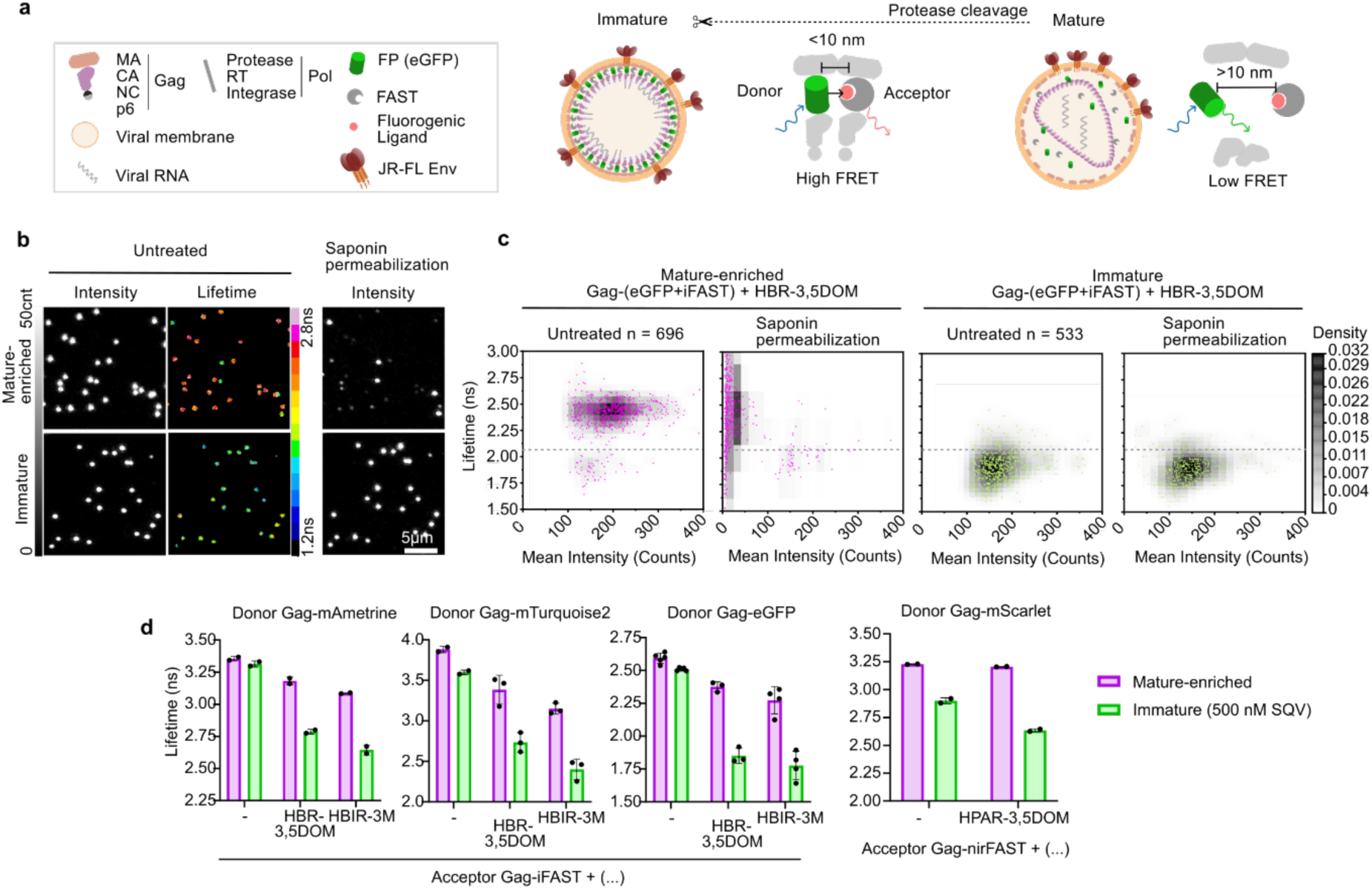
HIV-1 Gag processing detected by FRET–FLIM biosensor. **a.** Schematic of the FRET-based HIV-1 maturation biosensor. Donor and acceptor fluorophores inserted between the MA and CA domains of Gag are released upon protease cleavage, increasing their separation (>10 nm), thereby reducing FRET. **b.** Confocal images of mature-enriched (top) and immature (bottom) viral particles containing the Gag-(eGFP+iFAST) biosensor in the presence of the FAST ligand HBR-3,5DOM. Saponin addition triggers viral membrane permeabilization releasing fluorescent markers due to efficient Gag-processing. **c.** Density plots of fluorescence lifetime (𝜏DA) from individual particles showing distinct maturation-dependent populations before and after saponin treatment. **d.** Mean 𝜏DA values for seven donor-acceptor FRET combinations using donor fluorescent proteins (mAmetrine, mTurquoise2, eGFP or mScarlet) and acceptor FAST variants (iFAST or nirFAST) in absence (-) or presence of FAST ligands (HBR-3,5DOM, red emission; HBIR-3M, dark quencher or HPAR-3,5DOM, near infra-red emission). Each dot represents the mean 𝜏DA of at least 100 viral particles per experiment; bars show mean ± SD from independent experiments.

Gag processing can be indirectly detected as a loss of fluorescence once the viral membrane is permeabilized^32^. To evaluate whether the biosensor detects Gag processing during maturation, we measured FRET activity by FLIM and correlated lifetime values with fluorescent content loss upon viral membrane permeabilization (Fig. 1b,c). Mature-enriched samples (produced in absence of protease inhibitor) displayed an average donor lifetime in presence of acceptor (𝜏DA, Equation 1) of 2.39 ± 0.17 ns, whereas immature particles (+ 500 nM SQV) showed a reduced lifetime of 𝜏DA = 1.91 ± 0.14 ns, consistent with higher FRET in the immature state. Analysis of lifetime distributions from mature-enriched and immature Gag-(eGFP+iFAST) + HBR-3,5DOM (red emission fluorogen) biosensor samples (Supplementary Fig. 1) defined 𝜏DA = 2.1 ns as the threshold distinguishing mature from immature viral particles. Only 7.2% of mature-enriched particles fell below this threshold compared with 97.6% of immature particles. Membrane permeabilization of viral samples selectively released soluble fluorescent content from particles with 𝜏DA > 2.1 ns, confirming that lifetimes above the 2.1 ns threshold reflect efficient Gag processing.

We next tested the biosensor’s versatility to report Gag-processing using different FRET pair combinations (Fig. 1d and Supplementary Table 1). Immature particles consistently showed a decrease of 𝜏DA relative to mature-enriched particles (Δ0.4 ns to Δ0.75 ns depending on the FRET pair), directly linking biosensor response to viral maturation state. The various FRET pairs exhibit comparable FRET efficiencies (E = 52-63%, Equation 2, Supplementary Fig. 3a), and those incorporating eGFP or mTurquoise2 as donors coupled with iFAST–HBIR-3M (dark quencher) as the acceptor display a higher fraction of donor in interaction (fD, Equation 5) (Supplementary Fig. 3b). To evaluate the impact of biosensor incorporation on viral fitness, we compared the relative infectivity of each construct to WT, unlabelled virus and observed comparable values across viruses carrying different FRET-pair components (Supplementary Fig. 2). Among these, the pair Gag-(eGFP+iFAST) + iFAST–HBIR-3M exhibited the most favorable combination of infectivity and FRET performance (Supplementary Fig. 2 and Supplementary Fig. 3b, respectively) and was therefore used preferentially in subsequent experiments. Taken together, these findings establish the biosensor as a versatile platform for monitoring HIV-1 maturation, with optimized donor–acceptor pairs enabling multiplexed FLIM measurements and providing a basis for quantitative determination of viral maturation.

## Correlative FRET links early Gag processing with viral substructure analysis by cryo-ET

Relating early Gag processing events to structural changes in Env and the capsid formation is essential to understand the transitions determining viral fusion and infectivity. The clear separation between mature and immature viruses observed with Gag-(eGFP+iFAST) + iFAST–HBIR-3M biosensor at 37 °C (𝜏DA = 2.27 ns and 𝜏DA = 1.77 ns, respectively; Supplementary Fig. 2) suggested that the biosensor might also resolve intermediate maturation states. To link FRET-FLIM sensing of specific maturation states with their corresponding viral structures, we developed a correlative workflow enabling single-particle analysis. Viral particles incorporating the Gag-(eGFP+iFAST) + iFAST–HBIR-3M biosensor and the patient-derived JR-FL Env were vitrified ensuring optimal virus concentration to facilitate individual particle FRET-FLIM measurements (Fig. 2a). We avoided chemical fixation to preserve HIV-1 structures under near-native conditions. The level of Gag-processing was measured by FLIM in cryogenic conditions (Fig. 2b) and correlated to molecular details of individual virions (Fig. 2c). Tomographic analysis via cryoET provided morphological and spatial information of HIV-1 substructures to be classified according to maturation state (Fig. 2d).

**Fig. 2.**
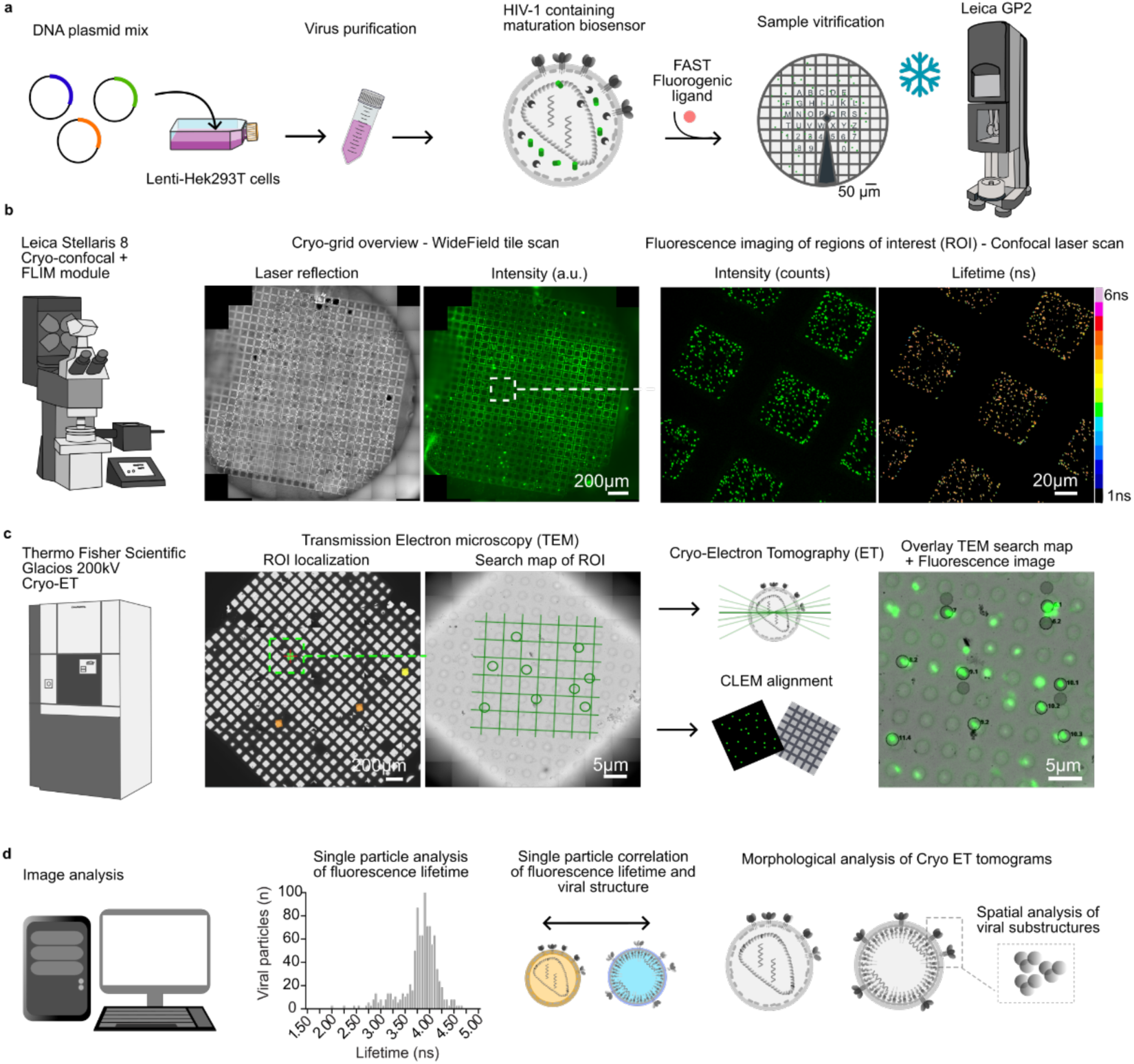
Lifetime-CLEM workflow for functional structure characterization of HIV-1. **a.** Schematic overview of sequential steps in Lifetime-CLEM workflow sample preparation. b. Widefield (left) and confocal (right) fluorescence images of cryo-grids containing viruses bearing the Gag-(eGFP+iFAST) biosensor. Regions of interest (ROIs; at least 10 per condition) showing homogeneous viral particle distribution were selected for subsequent FLIM imaging. c. Low magnification TEM images of cryo-grids enable ROI retrieval. Grid squares are then maps to identify holes in carbon support film (search map). CryoET data were then acquired at ∼10 holes per ROI. FLIM and TEM datasets were aligned (Overlay). d. Correlative single-virus analysis integrates lifetime and structural information, revealing FRET biosensor dynamics under cryogenic conditions in relation to HIV-1 maturation states, capsid morphologies, and Env distributions.

At cryogenic temperature, donor lifetimes in absence of acceptor (𝜏D) increased substantially in both mature-enriched samples (from 2.59 to 3.80 ns) and immature samples (from 2.51 to 3.78 ns; Fig. 3a,b and Supplementary Fig. 4a). Notably, adding the acceptor fluorogen expanded the FRET dynamic range from Δ0.5 ns at 37 °C to Δ1.03 ns at -195 °C. The fraction of interacting donor and FRET efficiency also increased, from 72.18% to 83.17% and from 57.88% to 69.11%, respectively (Supplementary Fig. 4b,c). These results show that the biosensor reports maturation-associated structural rearrangements under 37 °C and cryogenic conditions (−195 °C). We next investigated the correlation between fluorescence lifetimes observed under cryogenic conditions and individual viral particle structures visualized by cryo-ET (Fig. 3c). Individual particle FRET-FLIM analysis of mature-enriched samples predominantly exhibited τDA > 3.5 ns, corresponding to particles displaying HIV-1 capsids in cryo-ET, whereas immature samples (τDA < 3.5 ns) showed the typical immature Gag lattice^1^ covering >50% of the inner membrane surface. Samples treated with intermediate concentrations of SQV (25 nM) displayed a broader range of τDA values (2.4 ns - 4.5 ns) which correlated with different degrees of Gag lattice coverage or absence of internal structure. These particles may represent an intermediate stage of Gag processing, sufficient to disrupt the immature lattice to a certain extent but insufficient to complete capsid assembly. These observations demonstrate the power of functional correlative microscopy using FRET-FLIM biosensors to guide single-particle structural analysis and confirm that the biosensor can resolve intermediate states of HIV-1 maturation.

**Fig. 3.**
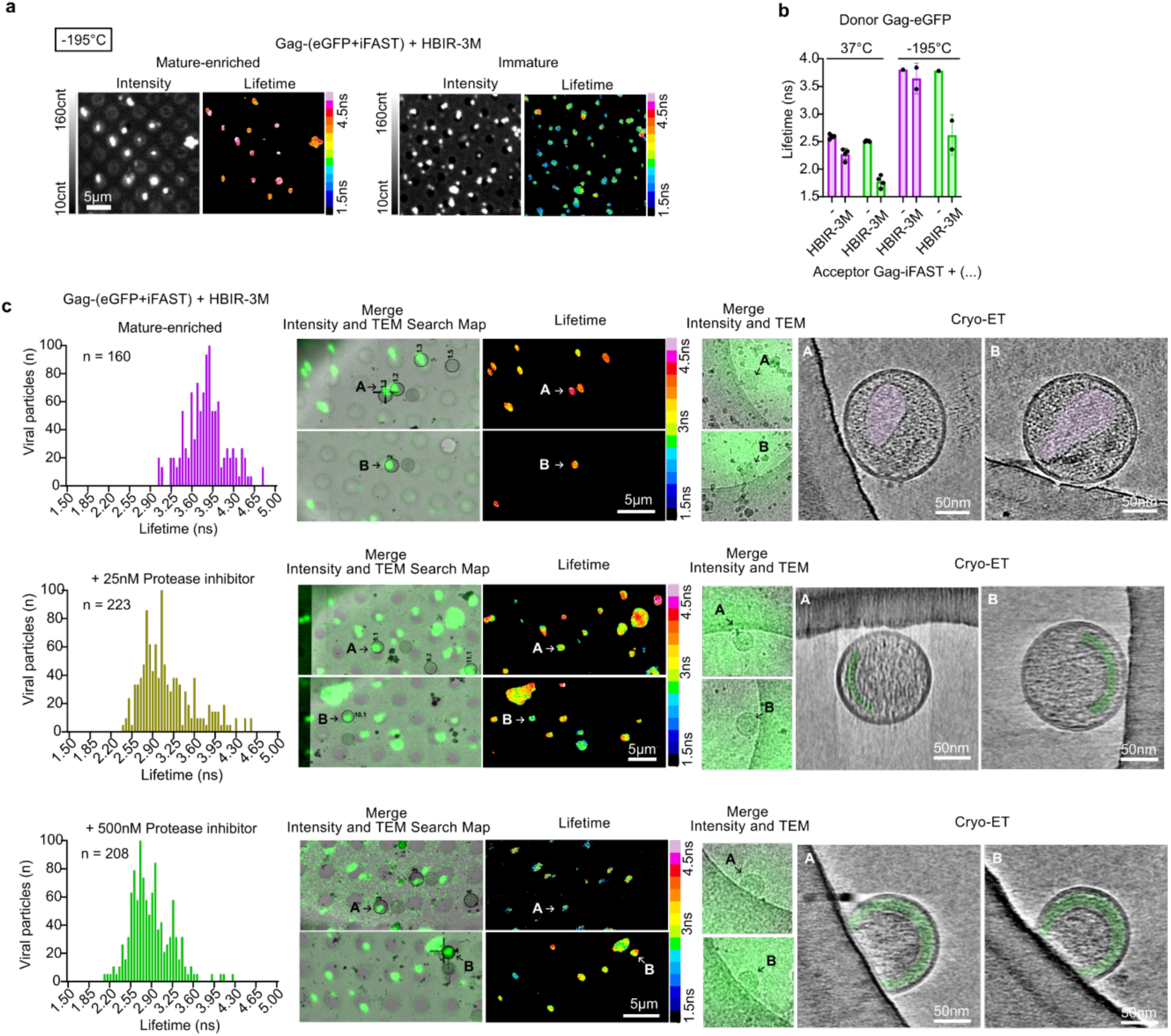
Lifetime-CLEM analysis of HIV-1 maturation states. **a.** Cryogenic FLIM images (–195 °C) showing fluorescence intensity (counts) and donor–acceptor lifetime (𝜏DA, ns) for mature-enriched (left) and immature (+500 nM SQV, right) viral preparations bearing the Gag-(eGFP+iFAST) + HBIR-3M biosensor. **b.** Comparison of mean 𝜏DA values from individual particles in mature-enriched (pink) and immature (green) samples imaged at 37 °C or -195 °C conditions. Each dot represents the mean 𝜏DA of at least 100 viral particles per experiment; bars show mean ± SD from independent experiments. **c.** Comparative Lifetime-CLEM analysis of mature-enriched and SQV-treated viruses (25 nM and 500 nM). Left: Frequency distributions of 𝜏DA values from at least 100 individual particles across ∼10 different FLIM ROIs at cryogenic temperature. Bars show the frequency distributions from 0.05 ns bins. Middle: Aligned FLIM intensity maps and TEM search images with lifetime information overlaid; arrows indicate representative particles (A, B) selected for cryo-ET. Right: Overlay of high-resolution TEM and fluorescence intensity of particles A and B, with corresponding representative cryo-ET slice from tomograms manually segmented to highlight mature capsids (pink) and immature Gag lattices (green).

## Morphological classification of HIV-1 core phenotypes by cryo-ET

Correlative analysis of viral samples produced in the presence of a suboptimal saquinavir concentration (25 nM) revealed an enrichment of particles exhibiting intermediate levels of Gag processing, characterized by an intermediate range of fluorescence lifetimes and reduced Gag-lattice coverage. To better characterize the frequency of this phenotype across the different viral samples, viral particles were classified according to their morphological phenotypes observed in tomograms (Fig. 4). In mature-enriched samples, the predominant morphology was conical capsid (77.19%), followed by tubular capsids (8.77%) and particles with multiple capsids (7.02%), reflecting known variability in HIV-1 core shapes^33,34^. Only 7.02% displayed an immature phenotype with >50% Gag lattice coverage of the inner viral membrane. Samples treated with 25 nM SQV showed immature phenotype with an overall reduced Gag lattice coverage; 33.85% showing >50% lattice coverage, 29.23% showed a partially detached lattice, 13.84% <50% coverage with some, 20% not showing a defined structure. Only 3.08 % of particles showed a conical capsid. Immature samples (500 nM SQV) showed a preserved Gag-lattice as 86.67% showed a >50% coverage, 4% showed <50% coverage and 9.33% displayed no visible structure. These results demonstrate that incorporation of our maturation biosensor minimally affects structural arrangements required for particle maturation. Reduced Gag lattice coverage observed in intermediate states of maturation indicates that Gag processing likely proceeds through sequential, locally initiated disassembly events rather than through a uniform, layer-by-layer dismantling of the immature lattice.

**Fig. 4.**
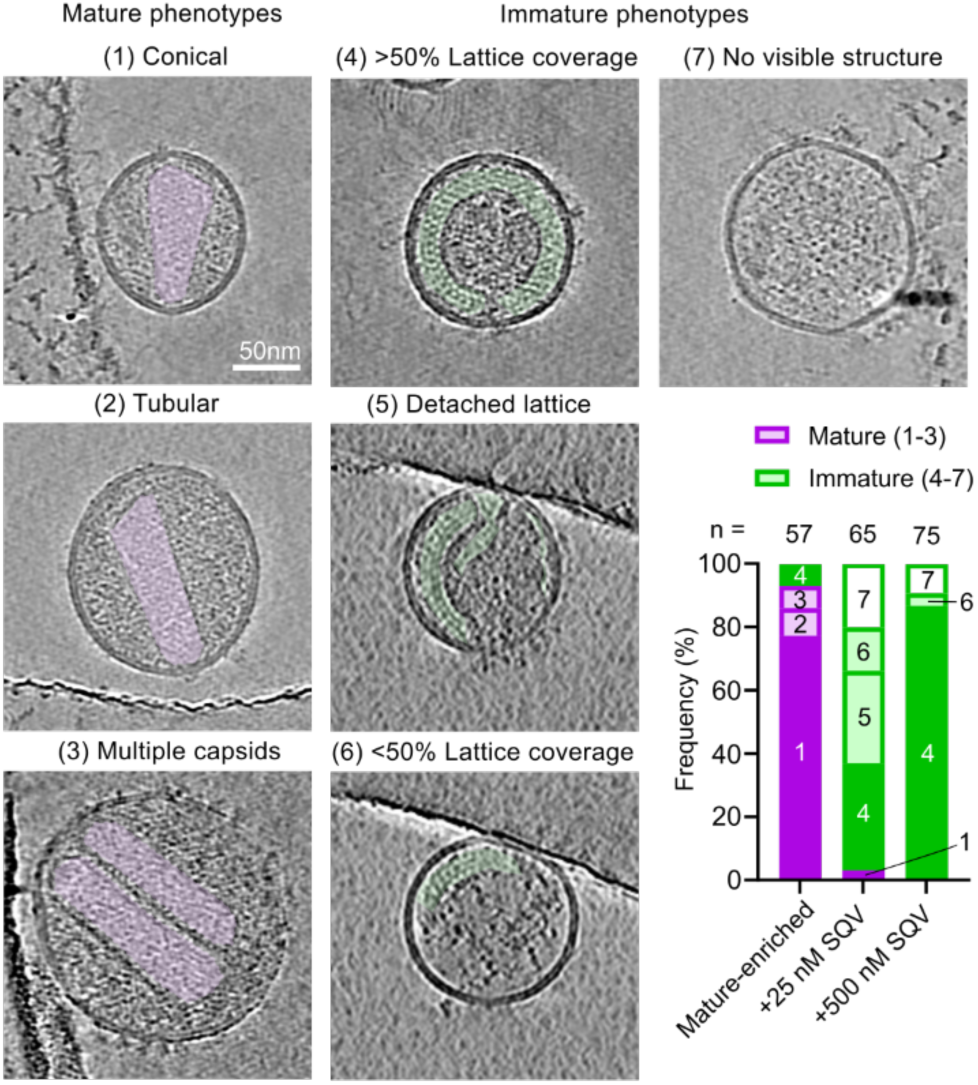
Cryo-ET analysis of HIV-1 maturation morphologies. Representative slice from cryo–electron tomograms of HIV-1 particles from mature-enriched, intermediate (25 nM SQV), and immature (500 nM SQV) samples. Manual segmentations highlight mature capsids (pink) and immature Gag lattices (green). Bar chart shows quantification of morphological phenotypes (% frequency) across the different viral preparations (n = number of total individual particles analyzed). Mature particles were classified into three capsid morphologies: conical (1), tubular (2), and multiple capsids (3). Immature particles were grouped into four categories based on lattice organization: >50% lattice coverage (4), partially detached lattice (5), <50% lattice coverage (6), and particles lacking visible structure (7).

## Spatial analysis of Env glycoprotein distribution on HIV-1 viruses at different maturation states

Our workflow preserves virion architecture without perturbing Env or core structures. We next examined how core organization across maturation states affects Env distribution. Tomograms were analysed to determine the spatial arrangement of patient-derived JR-FL Env trimers on the viral membrane (Fig. 5). Segmentation of the membrane enabled fitting of a three-dimensional hemispherical model of the viral surface, accounting for the missing-wedge effect. Coordinates of individual Env trimers were projected onto this model to calculate inter-trimer distances and examine their spatial distribution (Fig. 5). We evaluated whether JR-FL Env trimers were randomly distributed, dispersed or formed clusters relative to complete spatially random (CSR) simulations^35^, using Ripley’s function ^35^ and density ratios (Fig. 5b,c). In mature virions (n = 63), on average, Env trimers displayed higher H(r) values at sphere distances of 10–110 nm, indicating local clustering of Env proteins at the viral surface (Fig. 5c). At larger distances (230–290 nm), H(r) values fell below the CSR range, showing disperse distribution, resulting in an overall polarized distribution. In contrast, immature virions produced in the presence of 500 nM SQV (n = 49) exhibited H(r) values close to zero and within CSR confidence intervals across all distances, consistent with a random Env organization. Viruses produced with 25 nM SQV (n = 46) exhibited H(r) values at the edge of the CSR confidence interval, farther from zero than fully immature particles, suggesting the onset of cluster formation in virions at intermediate maturation states. Consistently, density-ratio analysis confirmed that mature virions exhibit local Env densities above the CSR confidence interval, with progressively lower densities in viruses treated with 25 nM SQV, approaching the random levels observed in fully immature particles (+500 nM SQV). Classification of individual particles according to the Env distribution profile by Ripley’s analysis (Fig. 5d) showed 53.06% of mature particles with a cluster Env distribution, 32.6% in intermediate viruses and 7.94% in immature particles. Nearest neighbour analysis showed median distance of 14 nm in mature virions (interquartile range 25^th^-75^th^ percentile, IQR: 7.47-22.84), 21.69 nm in intermediate (IQR: 11.02-38.57) and 22.81 nm (IQR: 13.44-36.11) in immature viruses (Supplementary Fig.5). Differences in Env distribution could not be attributed to Env numbers of viral particle size, as viral preparations exhibited comparable number of Env trimers: 13.39 ± 5.05 in mature particles, 8.86 ± 3.11 in intermediate virus or 9.86 ± 4.62 in immature particles (Fig. 5e) with an average virion size of radius = 70.52 ± 1.75 nm (Fig. 5f). These results show that release of matrix from Gag directly coordinates the structural reorganization of Env from a random to a clustered, polarized distribution at the viral surface, prior to capsid assembly.

**Fig. 5.**
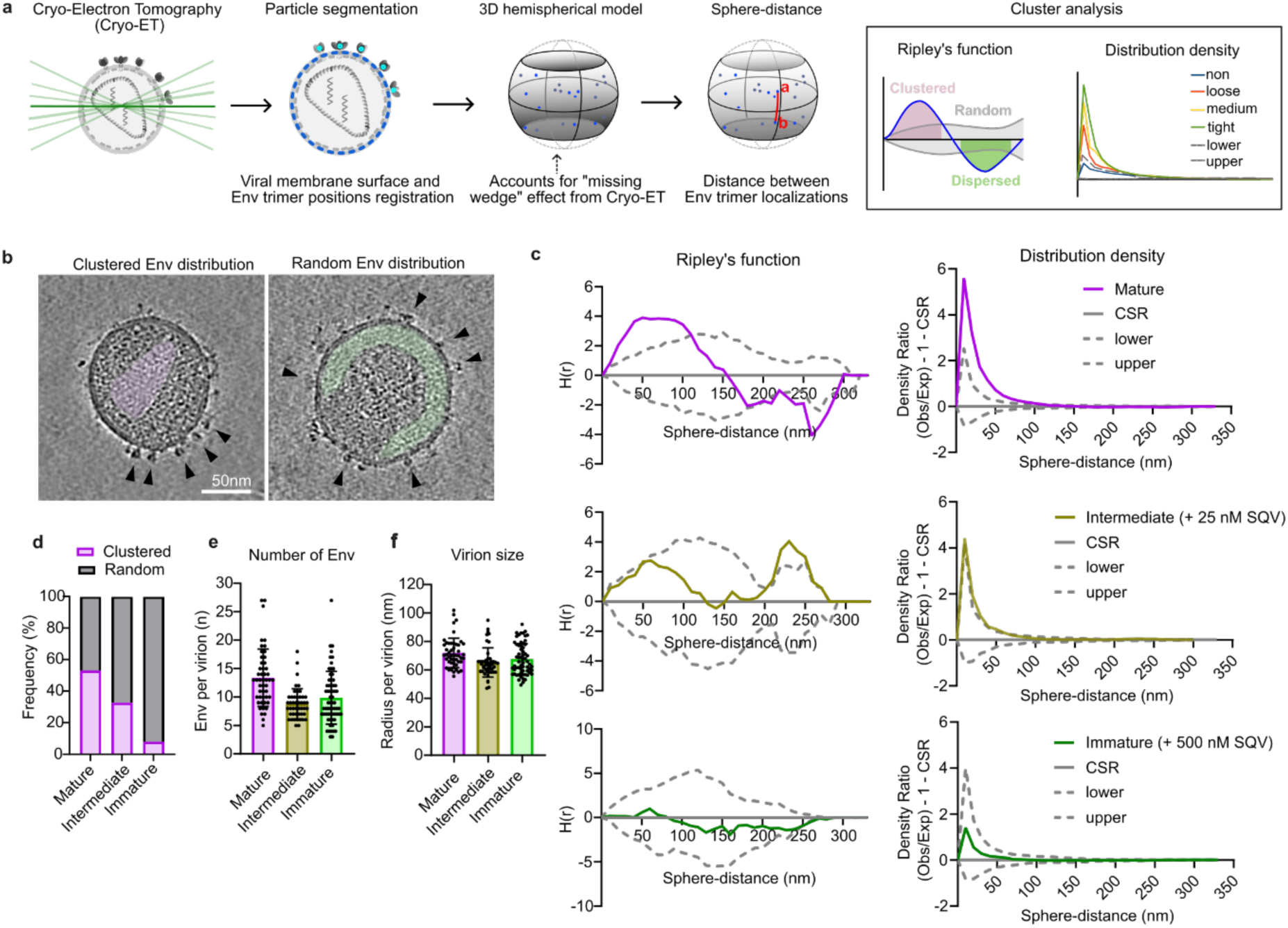
Maturation-dependent distribution of Env on HIV-1 surface. **a.** Schematic depiction of the analysis workflow for Env distribution from cryo-ET images. Viral particle segmentation and Env positioning allow reconstruction of a 3D hemispherical model of the virus to measure Env trimer spherical distances. Spatial analysis using Ripley’s functions and distribution density ratios provide information about Env clustering, dispersion or random distribution on the viral surface. **b.** Representative cryo-electron tomograms of a clustered Env surface distribution on mature particles (left) or random, homogeneous distribution on immature particles (right). Fully assembled, conical capsid is shown in pink. Immature lattice is highlighted in green. Arrows point at individual Env trimers on the viral membrane. **c.** Spatial distribution analysis using Ripley’s function H(r) (left) and density ratio from observed vs expected (random) events in a certain distance (right), from mature virions (top), virus treated with 25 nM SQV (middle) and 500 nM SQV (bottom). CSR: completely spatially random. Lower and upper: CSR 95% confidence intervals. **d.** Classification of individual viral particles as a function of Env distribution based on Ripley’s function H(r). **e.** Quantification of number of Env per viral particle. **f.** Quantification of radius per viral particle, per condition. **e. and f.** Dots show values per individual particle; bars show mean ± SD from all examined tomograms.

## Early HIV-1 maturation state correlates with fusion

HIV-1 maturation occurs extracellularly and leads to structural rearrangements that determine the outcome of the subsequent infection cycle. To assess the relationship between Gag-processing between the matrix and capsid subunits and early functional steps of viral entry, we produced viral samples in presence of increasing concentrations of protease inhibitor (5 nM - 500 nM SQV). We observed progressively higher FRET (lower lifetime values) with increasing inhibitor concentrations (Fig. 6a). Notably, intermediate inhibitor concentrations (25-50 nM) produced a broader distribution of single-particle lifetimes, similar to cryogenic conditions, indicating a heterogeneous population with variable Gag-processing efficiency.

**Fig. 6.**
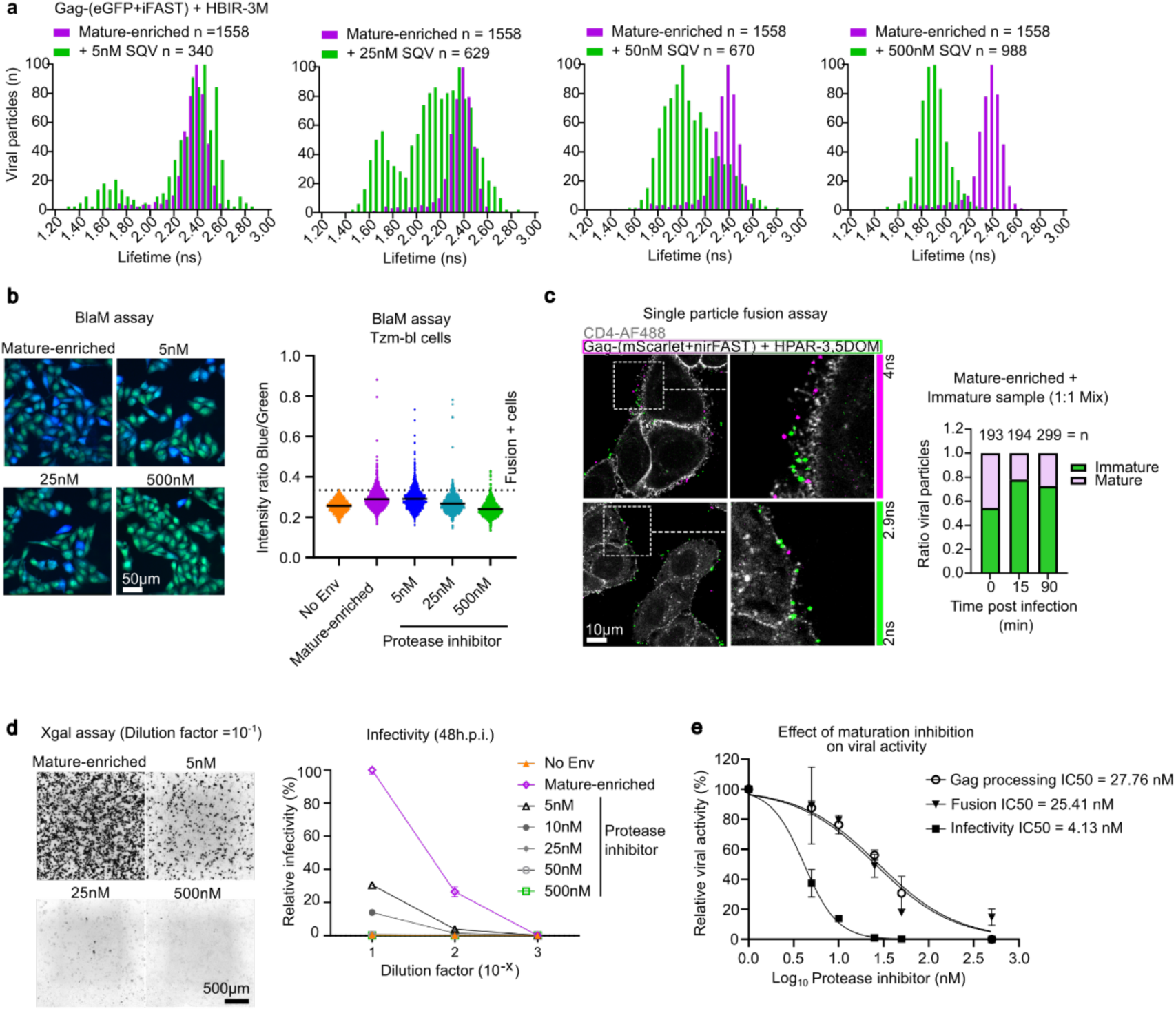
Functional implications of intermediate maturation states detected by the FRET-FLIM biosensor. **a.** Frequency distribution of 𝜏DA values from individual virions containing Gag-(eGFP+iFAST) + HBIR-3M biosensor produced in presence of increasing concentrations of protease inhibitor saquinavir (SQV), as indicated. Bars show the frequency distributions from 0.05 ns bins and are representative of three independent experiments. **b.** β-lactamase (BlaM) fusion assay in Tzm-bl cells with viruses bearing BlaM-Vpr produced in presence of increasing concentrations of SQV. Fusion efficiency was determined from the blue to green fluorescence emission ratio. Each dot represents a single cell. Values exceeding the maximum obtained from Env-deficient viruses (No Env = 0.334, dashed line) were considered fusion-positive events. **c.** Single-particle fusion assay using viruses bearing the Gag-(mScarlet+nirFAST) + HPAR-3,5DOM biosensor. A 1:1 mixture of mature-enriched and immature (+500 nM SQV) virus preparations was used to infect TZM-bl cells. Co-localization with immunostained CD4 (AF488) identified viral particles at the plasma membrane at 0, 15, and 90 min post-infection. Proportion of mature particles (𝜏DA > 2.9 ns, pink) and immature particles (𝜏DA < 2.9 ns, green) were quantified to determine fusion dynamics. **d.** Infectivity assay in Tzm-bl cells using viruses containing Gag-(eGFP+iFAST) produced at increasing concentrations of SQV. Infected cells were identified by high contrast coloration under transmission light microscopy. Relative infectivity was calculated from three sample dilutions and normalized to mature-enriched viral samples. **e.** Comparative analysis of maturation inhibition effects on Gag processing (n = 3), fusion (n = 2), and infectivity (n = 2). Nonlinear regression was applied to each dataset to determine IC50. Goodness of fit: Gag-processing, R^2^ = 0.9718; BlaM fusion, R^2^ = 0.8892; infectivity, R^2^ = 0.9917.

We next assessed the ability of viruses with distinct maturation states to fuse with the host plasma membrane and access the host cytosol. Viruses were produced incorporating the HIV-1 accessory protein Vpr fused to β-lactamase, and equal amounts of particles were used in a BlaM fusion assay^36^ on TZM-bl cells (Fig. 6b). At 90 min post-infection, single-cell analysis demonstrated a clear maturation-dependent modulation of fusion efficiency compared to viruses lacking Env (No Env). Viruses treated with 25 nM protease inhibitor yielded 48.94% fusion-positive cells relative to the mature-enriched sample, while fully immature viruses (500 nM SQV) displayed a pronounced decrease in fusion, with only 14.68% fusion-positive cells. To further confirm the dependence of efficient fusion on maturation, we performed a time-of-addition, single-particle experiment, applying an equal mix of mature-enriched and immature particles containing the biosensor to Tzm-bl cells (Fig. 6c and Supplementary Fig. 6). Gag(mScarlet+nirFAST) + HPAR-3,5-DOM (far red emission) was selected to facilitate multiplexing with immuno-stained CD4 with AF488. Individual particles were tracked for co-localization with the HIV-1 receptor CD4 at different time points (0, 15 and 90 min post infection) to assess entry. Discrimination between viral populations was defined by 𝜏DA = 2.9 ns based on the mScarlet biosensor range (Supplementary Fig. 1). At t = 0, 54.4% of analyzed particles displayed 𝜏DA < 2.9 ns increasing to 77.84% at 15 min and 72.58% at 90 min post infection, indicating that mature particles (𝜏DA > 2.9 ns) engage in fusion as early as 15 min post-infection.

To determine whether fusogenic particles with intermediate states of maturation remained capable to reach the nucleus and establish infection, we performed infectivity assays in TZM-bl cells (Fig. 6d). Infectivity was markedly reduced, dropping to 37.3% at 5 nM and 0.93% at 25 nM SQV, relative to mature-enriched particles, indicating that viral entry steps after fusion are highly sensitive to incomplete Gag processing, consistent with the lack of formed capsid observed in intermediate virus samples treated with 25 nM SQV (Fig. 4). Comparative analysis of FLIM-based Gag processing between MA/CA (Equation 6), fusion activity (BlaM assay), and infectivity revealed a high correlation between inhibition of matrix release from Gag and fusion (IC50 = 27.76 nM and 25.41 nM, respectively), whereas infectivity was ∼6x more severely affected (IC50 = 4.13 nM). These findings show that fusion competence is established early during maturation, when matrix release from Gag drives Env clustering, as detected by the biosensor. By contrast, full infectivity emerges only after subsequent maturation events culminating in complete capsid formation, revealing a temporal uncoupling between fusion activation and the acquisition of infection competency during maturation.

## Discussion

Here we have investigated how the coordinated structural transitions of the immature viral particle are coupled to the acquisition of host-interacting functions. Methods that directly link in-situ molecular events to structural biology remain limited. To address this gap, we have developed a novel FRET-FLIM-based approach that enables correlative analysis of dynamic molecular events with structural details using cryo-ET. We developed a modular biosensor that detects Gag-processing by changes in FRET. By optimizing fluorophore pairs, the sensor resolves transient intermediate states of maturation and supports multiplexing with additional fluorescent markers. Insertion of the biosensor between the MA and CA domains of Gag preserved virion functionality and efficiently reported release of MA from the Gag-lattice. Previous studies showed that cleavages downstream of CA are sufficient to rescue the impaired fusogenicity of immature virions^37^, and point mutations that alter MA lattice organization in mature virus do not reduce infectivity^22^. MA release reported by the biosensor coincided with increased fusion activity during early maturation. These observations indicate that MA release from the Gag lattice, rather than complete MA lattice maturation, is critical for initiating fusion. FRET-FLIM measurements guided the identification of intermediate states in cryo-ET with partially disassembled Gag lattices and absence of mature capsid. These intermediate states displayed a decoupling of fusion activation from productive infectivity, implying that CA structural maturation is not required for HIV-1 fusion. Premature viral entry during these states resulted in abortive infection, potentially due to exposure of uncoated viral genomes and consequent immune recognition^10^. Intermediate particles exhibited reduced lattice coverage, with local detachment from the membrane or absence of apparent structure, supporting a model in which Gag processing proceeds through local disassembly combined with displacive rearrangements of Gag-lattice sublayers, consistent with observations in protease cleavage mutants^21,38,39^. Genetic and structural studies have highlighted the importance of trimeric MA organization for Env cytoplasmic tail (CT) interaction, contributing to Env incorporation into virions^40,41^. Patient-derived JR-FL strain, unlike lab-adapted strains (ADA, BaL)^20,42,43^, possesses a full-length Env CT and incorporated relatively few Env trimers (∼5–25 per virion), consistent with prior observations^5,44^. Cryo-ET quantification may underestimate absolute Env numbers due to the “missing wedge” effect. Nevertheless, the relative differences in Env organization between experimental conditions remained similar. Previous work has shown that interactions between the Env CT and the immature Gag lattice restrict Env mobility, cluster formation and limit fusogenicity^4,5,37,45^. We demonstrate that early MA release coincides with the onset of Env clustering on the viral surface, correlating with enhanced fusion. Env polarization may compensate for low incorporation per particle, enhancing cooperative receptor engagement^43,46^. Maturation or cleavage of Env CT^47,48^ has been shown to induce exposure of conserved epitopes, suggesting that MA release from Gag allosterically modulates Env ectodomain conformation via the CT. Furthermore, surface clustering of Env may shield vulnerable epitopes, contributing to limit immune recognition^7^. Collectively, these findings highlight the importance of temporally coordinated Gag processing and MA release in regulating viral fusion, offering potential avenues for therapeutic intervention by targeting intermediate maturation states or disrupting MA–Env allosteric coupling.

Advances in correlative light and electron microscopy (CLEM) have introduced FRET detection in cells prior to sample vitrification^49,50^. However, pre-vitrification FRET remains limited by potential mismatch between the fluorescence signal and the final vitrified state, which reduce the fidelity of correlation with cryo-EM. In the context of HIV-1 biology, intensity-based detection of labelled virions combined with time-deterministic cryofixation has addressed these limitations by providing a precise temporal anchor to resolve chemically-induced viral uncoating^51^. Progress has also been made towards the integration of cryogenic super-resolution fluorescence microscopy to guide electron microscopy analyses^24,52^. However, super-resolution imaging only provides positional information and does not report on functional states or molecular interactions. Moreover, the altered photophysical and photochemical landscape at cryogenic temperatures further constrains the performance of these methods^24^. In contrast, the delayed photophysical processes of fluorophores at cryogenic temperatures—manifesting as prolonged excited-state lifetimes, enhanced photostability and quantum yield, and restricted dipole rotation^27,52–54^, —can be advantageous for FRET–FLIM. Cryogenic FRET–FLIM can facilitate the recovery of functional information by resolving conformational states or protein interactions occurring at <10 nm, within their native structural context for correlative cryo-ET analysis. Consistent with this, our biosensor displayed markedly longer fluorescence decays and increased FRET dynamic range at –195 °C compared with 37 °C, enabling identification of specific intermediate states of HIV-1 maturation. A key advantage of FLIM over intensity-based FRET measurements is that it relies solely on donor fluorescence to resolve FRET populations, making it inherently resistant to artifacts arising from spectral crosstalk and autofluorescence^55^. Working at cryogenic conditions introduces specific limitation such as contamination from water present in the environment forming ‘ice contamination’. This happens during transfer between instruments or to storage and during cooling in vapour phase during the FLIM acquisition. These issues may be mitigated by emerging contamination-reduction strategies in the cryo-EM field^56^.

Our approach leverages the sensitivity of FRET by FLIM to resolve HIV-1 maturation states at the single-particle level under both physiological and cryogenic conditions. This enabled us to correlate intermediate maturation states with their structural features by cryo-ET and to characterize their functional implications. We show that early matrix release from Gag reorganizes Env from a random to a polarized distribution before capsid assembly, with uncoupled functional consequences for fusion and infectivity. This insight highlights a potential window for antiviral intervention and informs vaccine design targeting HIV-1 maturation and entry. Our strategy permitted targeted interrogation of structural transitions initiated by a specific enzymatic cleavage and is readily adaptable to other biological systems.

## Methods Summary

Details of molecular cloning, light microscopy, FRET-FLIM, CryoFLIM, CryoET, Fusion and Infection assays are given in Methods.

## Online Content

Methods, along with any additional Extended Data display items and Source Data, are available in the online version of the paper; references unique to these sections appear only in the online paper.

## Data availability

Correlative cryo-FLIM and Cryo-ET, FLIM images will be deposited in the bioimaging Archive data repository (S-BIAD),

## Acknowledgments

This research was funded by the Chan Zuckerberg Initiative (CZI) “Multi-color single molecule tracking with lifetime imaging” (2023-321188) to S.S. and S.P.P., the European Research Council (ERC-2019-CoG-863869 FUSION) to S.P.-P., and Innovate UK Horizon Europe Guarantee “IMAGINE” (10048356) and Wellcome Career Development Award (225902/Z/22/Z) to M.G. This work was supported by the Wellcome Trust through the Electrifying Life Science grant (220526/Z/20/Z). The Rosalind Franklin Institute is funded by the UK Research and Innovation, Engineering and Physical Sciences Research Council. The electron microscope used here is supported by the Research complex at Harwell. Cryo-ET datasets were collected on the Glacios 2, using the Research Complex at Harwell at Diamond Light Source (proposal NT29493), and on the Titan Krios at the Rosalind Franklin Institute. The authors gratefully acknowledge the Microscopy Innovation Centre of King’s College London for their support and expertise in this work.

## Author Contributions

Conceptualization: I. C-A., D.J.W., S.P-P.; methodology and validation: I.C-A., D.J.W., V.G-G., R.T, A.L.B., P.N.W., M.D. and S.P-P.; formal analyses: I.C-A, D.J.W.,V.G-G. ; resources: A.G., M.G., M.D., and S.P-P.; data curation: I.C-A., V.G-G., D.J.W., T.S., R.T., A.L.B. and S.P-P.; writing & original draft preparation: I.C-A.; writing, review, and editing: I.C-A, V.G-G, R.T, A.L.B., P.N.W., M.D., M.G., S.S., D.J.W and S.P-P.; visualization: I.C-A, D.J.W., S.S. and S.P-P.; supervision: A.G., M.G., M.D., and S.P-P.; project administration: S.P-P.; funding acquisition: A.G., M.G., M.D., S.S. and S.P-P. All authors have read and agreed to the published version of the paper.

## ONLINE METHODS

### Cell culture

All cells were maintained in a humidified 5% CO2 atmosphere at 37°C. Virus producer cells, Lenti-X 293T (Takara Bio, 632180), were grown in phenol red-free Dulbecco’s Modified Eagle Medium/F12 (DMEM/F-12, Gibco) supplemented with 10% Fetal Bovine Serum (FBS, Thermo). Viral fusion and infectivity reporter cells, TZM-bl (NIH AIDS Research & Reference Reagent Program), were grown in high glucose DMEM (Thermo, 41965062) supplemented with 10% FBS. All cells were routinely mycoplasma tested (Eurofins genomics).

### Plasmid DNA

Plasmids pR8ΔEnv (containing the HIV-1 genome with a deletion in Env), pcRev (expressing HIV-1 Rev) and BlaM-Vpr (HIV-1 Vpr fused to β-lactamase) were kindly provided by Dr Greg Melikyan (Emory University, Atlanta, GA, USA). The pCAGGS plasmid containing JR-FL Env was a kind gift from Dr James Binley (Torrey Pines Institute for Molecular Studies, USA). Plasmids containing fluorescent reporter proteins were derived from pHIVGagiGFPΔEnv (bei resources, hrp-12455). Briefly, fluorescent proteins and FAST scaffold proteins were inserted between the matrix and capsid subunits of Gag, flanked by the HIV-1 protease recognition sites 5’-SQNYPIVQ-3’, using MluI and XbaI restriction enzyme sites (New England Biolabs, R3198 and R0145, respectively). Fluorescent protein sequences (i.e. mAmetrine, mTurquoise2, mScarlet) were amplified by PCR using Q5 polymerase (New England Biolabs, M0491) and mAmetrine-N1, mTurquoise2-N1 and pmScarlet_C1 (Addgene; 54505, 54843 and 85042, respectively) as template DNA plasmids. DNA template plasmids containing iFAST and nirFAST scaffold proteins were provided by Dr Arnaud Gautier (Sorbonne University). All plasmid sequences were verified by Nanopore sequencing (Genewiz).

### Pseudovirus production

Lenti-X 293T were seeded at a concentration of 5x10^5^ cells/mL in phenol red-free DMEM/F12 supplemented with 10%FBS. The following day cells were transfected using GeneJuice (Merck, 70967). DNA plasmid mix was prepared at a GeneJuice to DNA ratio of 3:1 in Opti-MEM Reduced Serum Medium (Gibco, 31985062). Biosensor-bearing pseudovirus were produced using pHIVGagFPΔEnv (being FP a fluorescent protein), pHIVGagFASTΔEnv (iFAST or nirFAST), pcRev and JR-FL, at a ratio of 2:2:1:3. Biosensor-containing virus are referred here after as Gag-(FP+FAST). For relative infectivity comparison assays, viruses containing unlabelled Gag were produced by partially (1:1) or fully (1:0) replacing labelled Gag plasmids with parental HIV-1 genome pR8ΔEnv. In case of fusion BlaM assays, DNA plasmid mix consist of pR8ΔEnv, pHIVGag-mTurquoise2ΔEnv, Vpr-BlaM, pcRev and JR-FL at a ratio of 10:1:10:2:6. In case of immature virus sample preparation, 500nM (unless indicated otherwise) of protease inhibitor Saquinavir mesylate (Sigma-Aldrich, S8451) was added onto cells on the day of transfection. Two days after transfection, virus containing media were harvested. Removal of cellular debris was achieved by centrifugation at 1000 x g for 10min at 4°C and further filtration through a 0.45µm filter. Virus samples were 200x concentrated by underlying a 10% Sucrose cushion and centrifugation at 10000 x g for 4h at 4°C. Viral pellets were gently resuspended in PBS, aliquoted and snap-frozen for long-term storage at -80°C.

### Infectivity assay

TZM-bl cells were seeded at a concentration of 2×10^5^ cells/mL in a flat-bottom 96-well plate. Serial dilutions of viral samples were prepared in complete DMEM and added onto cells. Two days after infection cells were fixed using 2% paraformaldehyde (PFA, Thermo 28908) for 15 min. Fixed cells were washed with PBS prior to adding pre-warmed X-gal solution (5 mM K3[Fe(CN)]6 (potassium ferricyanide), 5 mM K4[Fe(CN)]6 (potassium ferrocyanide), 2 mM MgCl2, 1 mg/ml X-Gal (5-bromo-4-chloro-3-indolyl-β-D-galactoside, Thermo 15520034), in PBS)) and incubated in the dark at 37 °C for minimum 2 h. Cells were imaged using transmitted light in a Leica DMi8 microscope (Leica Microsystems) equipped with a HC PL FLUOTAR 10x/0.30 DRY objective. Images were analysed using Fiji software. The number of infected cells was quantified from 2-3 technical replicates and a representative area of 2.2 mm^2^ was examined per replicate. A standard calibration curve was fitted to the results to calculate the concentration of infectious particles per µL (viral titre) using GraphPad Prism 10 software. The relative infectivity was calculated by normalizing the viral titre to the concentration of physical particles in the sample as assessed by confocal microscopy (see “quantification of viral particle concentration” section).

To calculate the IC50 of saquinavir protease inhibitor, data per condition was normalized by defining infectivity in the mature-enriched condition as 100%, and in 500 nM PI-treated viruses as the 0%. PI concentration values were transformed to base 10 logarithms. The results were analysed applying a non-linear fit log(inhibitor) vs. normalized response, with variable slope, using GraphPad Prism 10 software.

### Fusion assays BlaM assay

For β-lactamase fusion assay (BlaM assay), TZM-bl cells were seeded at a concentration of 1.5×10^5^ cells/mL in µ-Slide (Ibidi, 81816). The following day, equal numbers of physical particles per condition were diluted in cold Fluorobrite DMEM (Gibco, A1896701) supplemented with 2% FBS and 15mM HEPES (Thermo, 15630056). The concentration of physical viral particles resulting in a multiplicity of infection of 5 in the mature-enriched viral sample was considered as a reference to calculate the number of particles needed in all different conditions. Cells were washed with cold PBS prior to adding the viral inoculum and incubated for 1 h at 4 °C to allow virus adhesion onto the cell surface. Unbound viruses were washed and the media replaced with complete DMEM. Viral fusion was triggered by incubating cells for 90 min at 37°C 5% CO2. Media was replaced with Fluorobrite DMEM supplemented with 2% FBS and 15 mM HEPES and loaded with the β-lactamase substrate CCF2-AM from the LiveBLAzer FRET B/G Loading Kit (Thermo, K1095). Cells were incubated at RT in the dark for 2 h, washed with PBS and fixed with 4% PFA for 15 min at RT prior to imaging. Cells were imaged using a Wide-Field Leica DMi8 microscope (Leica Microsystems).

### Single particle fusion assay

For single particle fusion assay, TZM-bl cells were seeded at a concentration of 10^5^ cells/mL in a µ-Slide (Ibidi, 81816). The following day, a mixed sample of mature-enriched and immature viruses bearing the Gag-(mScarlet+nirFAST) biosensor was prepared at a ratio of 1:1, so that a total of 150 viral particles/cell, per condition, were diluted in cold Fluorobrite DMEM (supplemented with 2% FBS and 15mM HEPES) and added onto cells. Viral adhesion to cells was facilitated by spinoculation at 2000 x g for 20 min at 4 °C. Unbound virus was removed and samples were either fixed (t = 0) using 4% PFA or incubated in presence of complete DMEM for further 15 min or 90 min at 37 °C 5% CO2, followed by fixation. Fixed cells were immunostained to localize the HIV-1 receptor CD4 on the cellular membrane using a human anti-CD4 antibody coupled to AlexaFluor 488 (clone OKT4, BioLegend, 317419) at 0.5 µg/mL concentration. Fluorogen HPAR-3,5DOM was added to the samples immediately prior to imaging using a Stellaris confocal microscope (Leica Microsystems).

### Fluorogenic labelling of FAST biosensor viruses

Labelling of viral particles containing the FAST biosensor was performed by adding 5µM concentration of the appropriate fluorogenic chromophore diluted in PBS to the sample, immediately prior to image acquisition or plunge freezing (in the case of cryo-EM sample preparation). Ligands used for iFAST are the dark chromophore HBIR-3M ((4-hydroxy-3-methylbenzylidene)-4-thioxothiazolidin-2-one) and fluorogen HBR-3,5DOM (4-hydroxy-3,5-dimethoxybenzylidene rhodanine), whereas the fluorogen for near-infrared FAST (nirFAST2.0) is HPAR-3,5DOM (4-hydroxy-3,5-dimethoxyphenylallylidene rhodamine)^29,30^. FAST ligands were kindly provided by Dr Arnaud Gautier (Sorbonne University) and are also available from the Twinkle Factory (HBIR-3M a.k.a. ^TF^Darth, 514000-250; HBR-3,5DOM a.k.a. ^TF^Coral, 516600-250 and HPAR-3,5DOM a.k.a. ^TF^Carmine, 636715-250).

### Fluorescence lifetime imaging

Fluorescence lifetime imaging of virions on glass surface was performed using the MicroTime200 (Picoquant) time resolved fluorescence microscope. The incubator chamber (DigitalPixel) was preheated to 37°C. The viral sample was excited using diode lasers (440nm for Gag-mAmetrine and mTurquoise2 labelled viruses and 485 nm for Gag-GFP) (LDH series Picoquant) with a repetition rate of 20 MHz. The laser beam was coupled to an Olympus IX73 inverted microscope and focused onto the sample by a 60x Water, NA 1.2 (Olympus UPLSAPO). Emission passed through a quad-dichroic mirror (at 440, 485, 594, and 635 nm, Chroma) and a 100 μm pinhole (ThorLabs). The remaining emission was separated at a 560LP dichroic (Chroma). Wavelength emission for mAmetrine, mTurquoise2 and GFP was passed through a 525/50B filter (Chroma) before single-photon detection via a PMA hybrid detector (Picoquant). Time correlated single photon counting (TCSPC) was performed using the Multiharp 150 (Picoquant) and lifetime information was registered with Symphotime software (Picoquant).

Fluorescence lifetime of viral samples bearing Gag-mScarlet was performed using an inverted Stellaris 8 Falcon time resolved confocal microscope (Leica microsystems). Gag-mScarlet labelled viruses and CD4 stained with AF488 were excited using a pulsed white-light laser tunned at 561 nm and 488 nm, respectively, with a 20 MHz repetition rate. The laser beam was focused through a HC PL APO CS2 63x Oil NA 1.4 objective and out-of-focus emission was filtered through a 95.5 µm pinhole. Detection range was set up between 500-550nm for AF488 emission and 570-620nm for mScarlet. Emission was collected using HyD X detectors in photon counting mode. Sequential scanning was performed stack by stack with a frame accumulation of 50 and 120 nm pixel size.

Cryogenic fluorescence lifetime imaging microscopy (cryo-FLIM) was performed on an upright Stellaris 8 Falcon microscope (Leica Microsystems) equipped with a cryogenic stage and cryogenic HC PL APO 50x/0.90 DRY objective. Samples were imaged under cryogenic conditions (−194 °C). GFP marker was excited with a pulsed white-light laser tunned at 488 nm and 20 MHz repetition rate. Emission was filtered through a 247.5 µm pinhole and collected between 500-550 nm using a hybrid detector (HyD S2) in photon counting mode. Laser scanning was performed with a frame accumulation of 70 and 108 nm pixel size.

### Wide-Field imaging

TZM-bl cells loaded with CCF2-AM were imaged using Wide-Field Leica DMi8 microscope (Leica Microsystems). The sample was sequentially illuminated with transmission light and a continuous laser at 405 nm to excite CCF2-AM. Excitation light was focused through a HC PL FLUOTAR 20x/0.50 DRY objective. Emitted fluorescence was bifurcated using dual emission image splitter OptoSplit II (Cairn Research Ltd) incorporating a long pass dichroic mirror filter at 510 nm and emission filters ET480/40m and ET525LP (Chroma). “Blue” emission between 460-500 nm correspond to the donor fluorescence and “Green” emission >525 nm to the acceptor fluorescence. A neutral density filter of 32% transmittance at 546 nm was used to attenuate the >525 nm fluorescence emission. Emission was detected using a Prime 95B Scientific CMOS (sCMOS) camera (Teledyne photometrics) set at 10ms exposure time for transmitted light detection and 300 ms exposure time for fluorescence emission detection. At least 10 images of 659.58 µm x 329.79 µm size were obtained per channel from 2 technical replicates.

### Quantification of viral particle concentration

Serial dilutions of viral samples were prepared in PBS and placed onto glass bottom µ-Slides (Ibidi, 81817). Samples were incubated at room temperature for at least 1 h to allow the particles to adhere onto the glass surface. Images were acquired using a MicroTime200 confocal microscope (Picoquant). At least 3 representative areas of 60µm^2^ were analysed per condition and the number of physical particles quantified using Fiji software. A standard calibration curve was fitted to the results to calculate the concentration of viral particles per µL using GraphPad Prism 10 software.

### Fluorescence lifetime analysis

The average lifetime non-fitting approach allows calculation of the average lifetime per pixel without applying any mathematical model. All time correlated single photon counting (TCSPC) measurements were recovered as PicoQuant PTU files with Symphotime software (Picoquant) or LAS X microscope software (Leica microsystems). Multi-frame images corresponding to time-correlated fluorescence intensity were extracted using PTU_Reader v.0.0.9 plugin for ImageJ (https://github.com/UU-cellbiology/PTU_Reader). Home-made software was developed in ImageJ to calculate the total amount of photons and average lifetime per pixel from time-correlated multi-frame intensity images, applying the following formula:

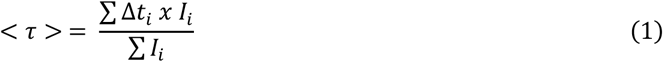

Where Δ𝑡_!_ is a particular delay time relative to the laser pulse and 𝐼_!_ the corresponding intensity in that particular time frame. ∑ 𝐼_!_ corresponds to the total amount of photons in all frames. Intensity frames corresponding to the raise of the laser pulse were excluded from the analysis so that t = 0 was set at the start of the fluorescence intensity decay. A cut-off intensity of 50 photons and an area of 5-100 pixels/particle was considered to analyse the average lifetime per individual viral particle. Frequency distributions of average lifetime values per viral particle were calculated and normalized by defining the largest value in each data set as 100%. A nonlinear regression Gaussian fit was applied to the resulting frequency distributions using GraphPad Prism 10 software.

### FRET and binding efficiency analysis

The FRET efficiency (E) from lifetime measurements was calculated applying the following formula:

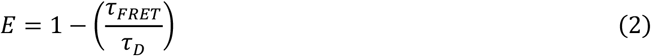

Where 𝜏*_FRET_* is the lifetime of the donor population engaged in FRET and 𝜏*_D_* the lifetime of non-interacting donor. In our system, immature HIV-1 particles produced in presence of protease inhibitor (500nM SQV) preserve the biosensor in the highest FRET regime conformation. Therefore, 𝜏*_D_* was obtained by applying a mono-exponential fit to the fluorescence decay from immature viruses in absence of acceptor, as follows:

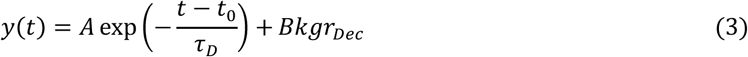

Where *A* is the pre-exponential factor (“amplitude”) and 𝐵𝑘𝑔𝑟*_Dec_* is the correction for background (afterpulsing, dark counts, environmental light).

𝜏*_FRET_* was calculated by applying a double-exponential fit to the fluorescence decay curve from an immature sample in presence of acceptor, and fixing the parameter 𝜏*_D_* to the value obtained in Eq. 3, using the following formula:

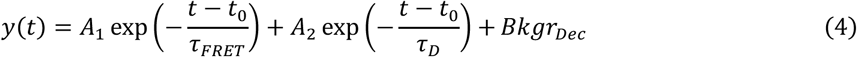

Where 𝐴_1_ is the pre-exponential factor of 𝜏*_FRET_* (proportion of donor engaged in FRET) and 𝐴_2_ the pre-exponential factor of 𝜏*_D_* (proportion of non-interacting donor).

Binding efficiency of FRET pairs was calculated as the fraction of interacting donor:

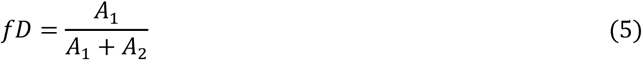

Decays were analysed using SymphoTime 64 software (Picoquant) and FLIMJ^57^ plugin in Fiji.

### Gag-processing efficiency analysis

Our HIV-1 maturation biosensor reports changes in donor lifetime in presence of acceptor as a function of Gag processing by the HIV-1 protease. To calculate the extent of Gag-processing in viral samples produced in the presence of different concentrations of protease inhibitor (PI) we applied the following formula:

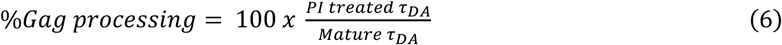

Where 𝑃𝐼 𝑡𝑟𝑒𝑎𝑡𝑒𝑑 𝜏*_DA_* is the mean average lifetime of viral particles produced in presence of protease inhibitor and 𝑀𝑎𝑡𝑢𝑟𝑒 𝜏*_DA_* is the mean average lifetime of viral particles produced in absence of protease inhibitor. At least 500 viral particles were analysed per condition. Results were normalized by defining the %Gag-processing in mature-enriched condition as 100%, and in 500 nM PI-treated viruses as the 0%. PI concentration values were transformed to base 10 logarithms. To calculate the IC50 of Saquinavir protease inhibitor, data were fit to a non-linear fit log(inhibitor) vs. normalized response, with variable slope, using GraphPad Prism 10 software.

### Fusion efficiency analysis

Data from the HIV-1 fusion BlaM assay were recovered using LASX software (Leica microsystems) and analysed using home-made scripts (ImageJ, https://imagej.net/ij/), including single cell segmentation, fluorescence quantification per channel and ratiometric calculation of “Blue” (460-500 nm) to “Green” (>525nm) signal for each individual cell. Fusion-positive cells were considered when Blue to Green ratio from CCF2-AM signal was higher than the maximum value obtained in the “No Env” condition (viruses deployed of Env protein). Percentage of fusion-positive cells relative to the total cells was calculated per condition from at least 500 cells. To calculate the IC50 of saquinavir protease inhibitor, data were normalized by defining fusion in the mature-enriched condition as 100%. PI concentration values were transformed to base 10 logarithms. The results were adjusted to a non-linear fit log(inhibitor) vs. normalized response, using GraphPad Prism 10 software.

For single-particle fusion analysis using Gag-mScarlet viruses, cut-off intensity of 50 photons and a particle size of 5-100 pixels were considered to calculate the average lifetime per viral particle. Only particles co-localizing with the CD4 marker were considered for the analysis. Viruses were classified as mature if average lifetime was above 2.9 ns and immature when average lifetime was equal or below 2.9 ns. Percentages of each type of viruses were calculated per condition.

### Cryo-EM Sample Preparation

Purified viral particles were diluted in PBS resulting in a final concentration of ∼10^6^ particles/µL in presence or absence of 5 µM HBIR-3M fluorogen. For plunge freezing, 2 µL of sample was applied to glow-discharged (20 mA, 30 s) holey carbon grids (gold R2/2, 300 mesh, finder type, Quantifoil, wash protocol 3). Grids were blotted for 4 s using a Leica GP2 (Leica Microsystems) operated at 4 °C and 70 % humidity and immediately plunged into liquid ethane cooled by liquid nitrogen. Grids were clipped and stored under liquid nitrogen until imaging.

### Tilt Series acquisition

Tilt series were acquired on a Glacios 2 (Thermo Fisher Scientific) operating at 200 kV, or a Titan Krios (Thermo Fisher Scientific) operating at 300 kV, both equipped with a Falcon 4i direct electron detector and a Selectris post column (10 eV) energy filter on the Titan Krios. Low magnification grid square overview images were acquired at a magnification of 470x and search maps were acquired at an intermediate magnification of 8700x (corresponding to a pixel size of 33.94 Å/pix). For tilt series acquisition, all images were recorded in counting mode and saved in eer format, with an average dose of 3.2 e-/Å^2^/tilt. Tilt series were collected using Tomo5 (Thermo Fisher Scientific) at a magnification of 81000x (corresponding to a pixel size of 1.95 Å/pix on Glacios 2 and 1.57 Å/pix on Titan Krios) and a target defocus of -3 to -5 μm. The data collection scheme was grouped dose-symmetric schemes spanning ± 60° in 3° increments, with total dose limited to ∼ 132 e-/Å^2^.

### Tomograms reconstruction

For data acquired using the Glacios 2, tomograms were reconstructed either using the automated tomography pipeline implemented at the UK national electron Bio-Imaging Center (eBIC)^58^ at Diamond Light Source, or manually using IMOD^59^. For the automated reconstruction, motion correction was performed with MotionCor2^60^ and CTF estimation using CTFFind4^61^. Frames were then aligned and tomograms reconstructed using Aretomo^62^ using the SART like reconstruction and denoised using Topaz^63^ for visualization. For manually reconstructed tomograms, tilt-series were pre-processed using Warp^64^ for motion correction and CTF estimation followed by reconstruction in IMOD using weighted back-projection, with a simultaneous iterative reconstruction technique (SIRT)-like filter^65^ applied for visualization. For data acquired on the Titan Krios, tilt series were pre-processed similarly in Warp^64^, followed by reconstruction in Aretomo^62^ and filtered with IsoNet^66^.

### CLEM alignment

To align cryo-electron tomography (cryo-ET) search maps with cryogenic confocal microscopy images, we used the BigWarp plugin in Fiji. Virus particles and hole patterns served as fiducial markers, and ≥4 corresponding points were manually identified in both modalities. The selection of fiducial points was guided by the orientation of the finder grid, enabling accurate alignment between atlas views, and an affine transformation was computed to register the confocal image stack to the cryo-ET coordinate system.

### Env distribution analysis

Viral particles from reconstructed tomograms were subjected to analysis to obtain Env locations and calculate inter-trimer distances. Individual Env trimer positions and viral membrane boundaries were manually picked using IMOD^59^. Average distances of segmented viral particles were used to model an hemisphere accounting for the cryo-ET “missing wedge” effect. Env trimer positions were projected onto the modelled hemisphere and point-pattern analysis was performed using nearest neighbour distance, density analysis and Ripley’s functions as described previously^35^, to obtain information about Env level of clustering. Analyses were performed using R and plotted using GraphPad Prism 10 software.

