## Supplementary Information for "Early HIV-1 maturation drives Env clustering and fusion competence"

*Corresponding authors at:

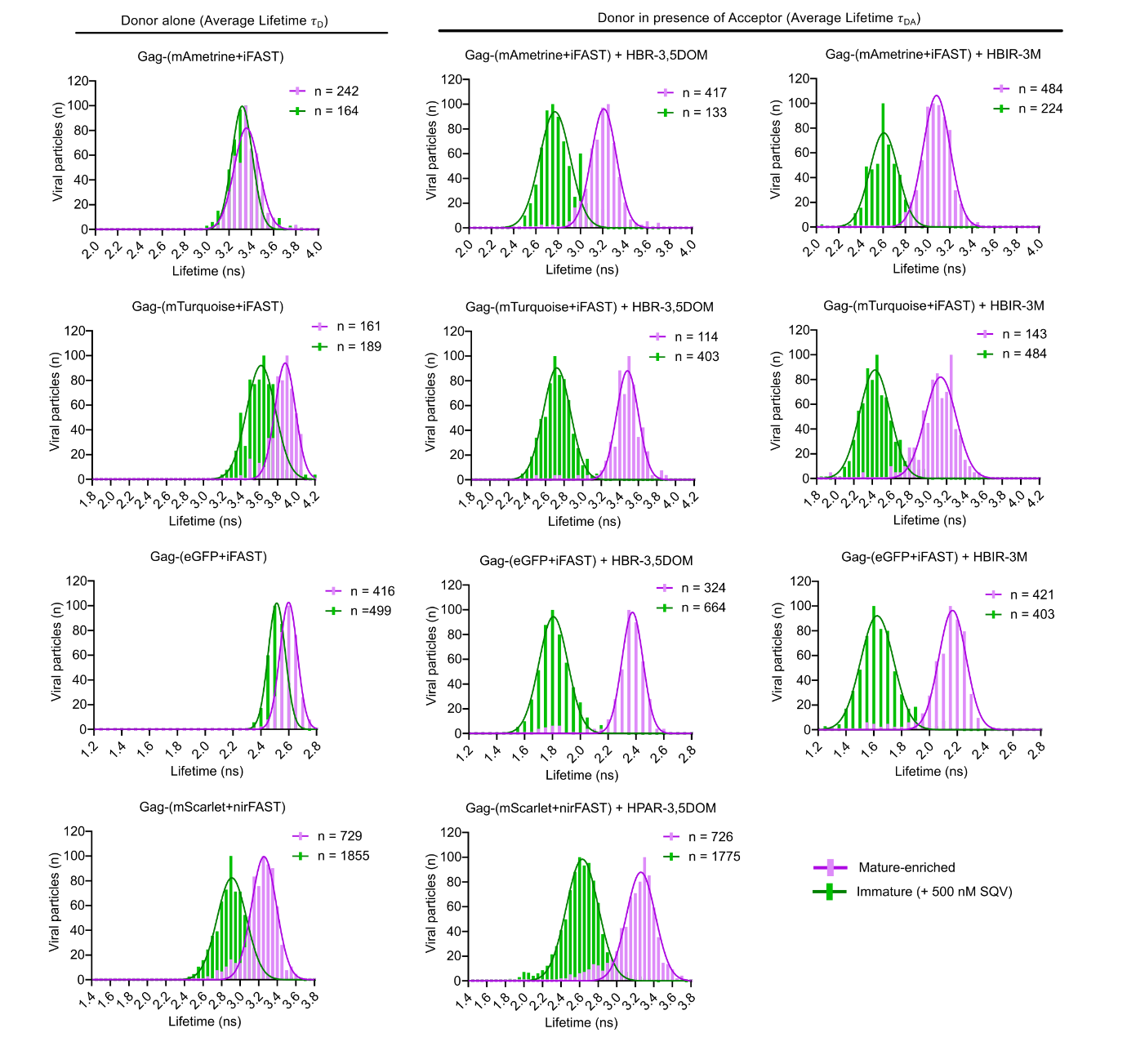

**Supplementary Figure 1. Biosensor response to protease inhibition.** Frequency distribution of average lifetime values from individual virions containing seven different combinations of the FRET-FLIM biosensor produced in absence (mature-enriched, pink) or presence of protease inhibitor (immature, + 500 nM SQV, green). Left: ditribution of average lifetime values in absence of acceptor (𝜏D). Right: ditribution of average lifetime values in presence of acceptor (𝜏DA). Number of individual particles analyzed is indicated for each condition and it is representative of at least two independent repeats. Bars show the frequency distributions from 0.05 ns bins and a nonlinear regression Gaussian fit was applied to the resulting data.

**Supplementary Table 1. Properties of donor and acceptor fluorophores.**

| FRET pair component | Name | λ_abs_ (nm) | λ_em_  (nm) | Visible colour | EC (mM^-1^ cm^-1^) | QY | Brightness | Lifetime (ns)  in live cells | Ref. |
| --- | --- | --- | --- | --- | --- | --- | --- | --- | --- |
| Donor | mAmetrine | 406 | 526 | green | 45 | 0.58 | 26.1 | 3.5 | ^1,2^ |
|  | mTurquoise2 | 434 | 474 | blue/  green | 30 | 0.93 | 27.9 | 4.1 | ^2,3^ |
|  | eGFP | 488 | 507 | green | 55.9 | 0.6 | 33.54 | 2.7 | ^2,4,5^ |
|  | mScarlet | 569 | 594 | red | 100 | 0.7 | 70 | 3.4 | ^2,6^ |
| Acceptor | iFAST +  HBIR-3M | 514* | 567* | dark | 12* | 0.003* | 0.036* | N/A | ^7^ |
|  | iFAST +  HBR 3,5 DOM | 519 | 600 | red | 44 | 0.33 | 14.52 | 2.48 | ^7^ |
|  | nirFAST + HPAR-3,5DOM | 636 | 713 | far red | 46 | 0.067 | 2.278 | n.d | ^8^ |

Abbreviations are as follows: λ_abs_ wavelength of maximal absorption; λ_em_ wavelength of maximal emission; EC, extinction coefficient at λ_abs_; QY, fluorescence quantum yield; Brightness = QY x EC; N/A, non-applicable; n.d. non-defined. *Reference numbers reported for pFAST + HBIR-3M^7^.

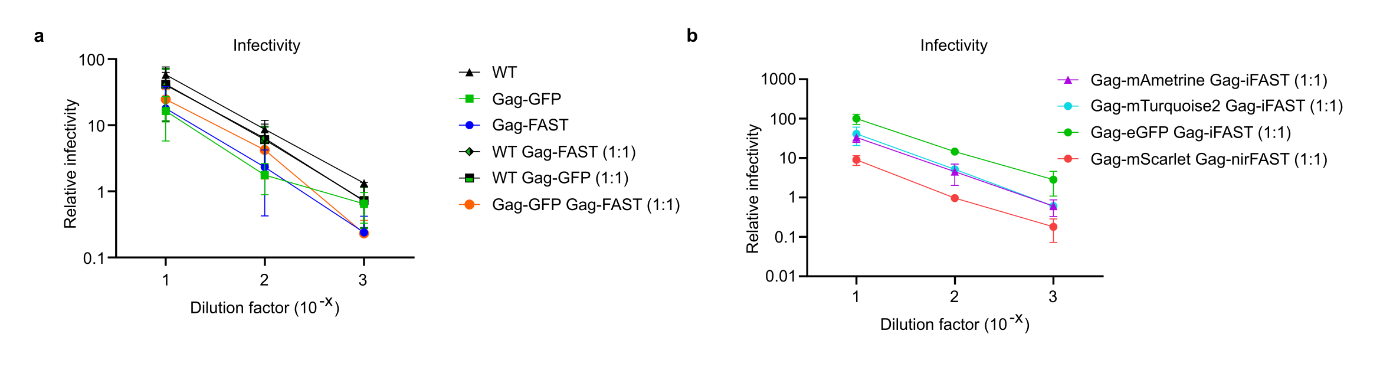

**Supplementary Figure 2. Infectivity assays of viruses containing the FRET-FLIM biosensor. a.** Comparative infectivity analysis of unlabelled wild-type (WT) viruses *vs* fully Gag-labelled viruses incorporating GFP, FAST, an equal mixture of GFP and FAST (1:1), or partially labelled viruses containing either GFP or FAST in a 1:1 ratio with unlabelled Gag. **b.** Comparative infectivity of viruses carrying the FRET-FLIM biosensor with different fluorescent proteins as donors and variants of FAST as acceptors in a 1:1 ratio. **a,b.** Tzm-bl were infected with 10-fold serial dilutions of virus preparations containing equal numbers of physical particles, and relative infectivity was quantified by X-gal staining and normalized to WT (**a**) or GFP (**b**) levels.

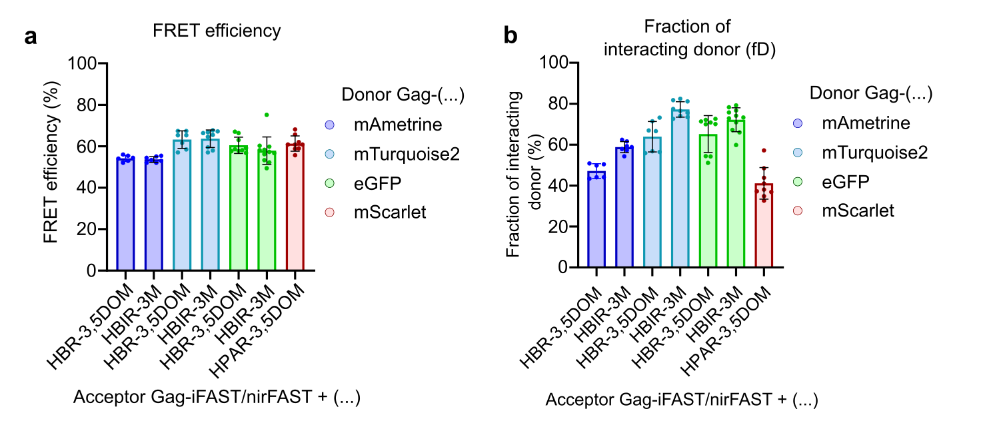
**Supplementary Figure 3. FRET efficiency of the biosensor pairs in virions.** Bars show the mean ± SD of **a.** FRET efficiency or **b.** Fraction of interacting donor, obtained from double-exponential fit analyses of the donor in presence of acceptor (𝜏FRET) and donor alone (𝜏DA). Each dot represents a double-exponential fit from a single FLIM image.

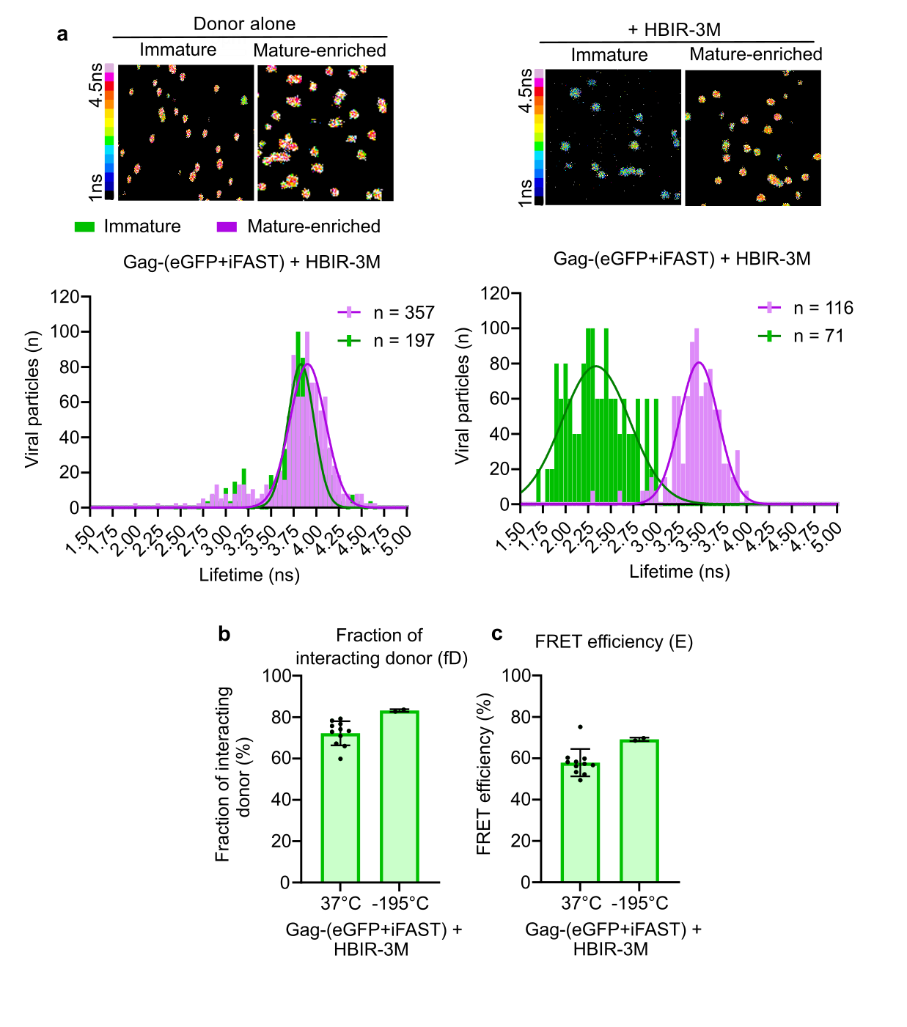

**Supplementary Figure 4. Cryo-FLIM measurements of viral particles containing the maturation biosensor. a.** Cryo-confocal FLIM images (upper row) of virions on cryo-grids containing Gag-(eGFP+iFAST) with or without HBIR-3M as an acceptor. Frequency distribution of average lifetime values from individual virions. Bars show the frequency distributions from 0.05 ns bins from mature-enriched samples (pink) or immature samples (+ 500 nM SQV, green) and a nonlinear regression Gaussian fit was applied to the resulting data. **b.** Fraction of interacting donor and FRET efficiencies in % of the eGFP biosensor at 37 °C or -195 °C. Bars represent the mean ± SD of fits from FLIM images. Each dot represent the fit from an individual FLIM image.

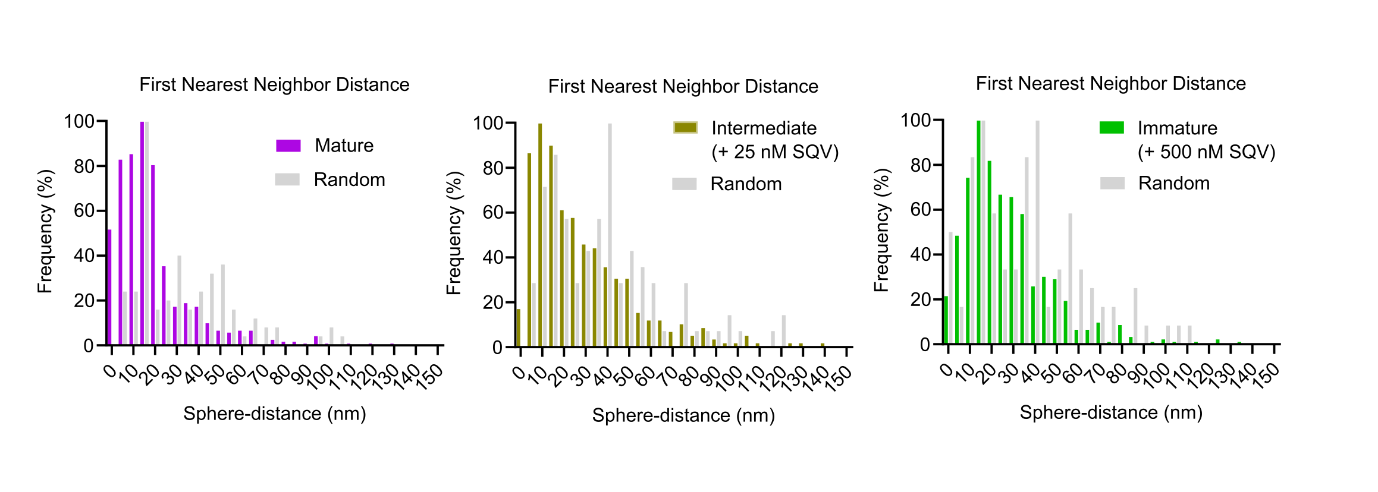

**Supplementary Figure 5. Distribution of Env distances.** Each Env position on HIV-1 virions was assessed for the distance to the nearest neighbor Env on the viral membrane of Mature (pink), Intermediate (yellow; from samples treated with 25 nM SQV) and Immature particles (green; from samples treated with 500 nM SQV). Grey bars represents data from 100 HIV-1 random Env distribution simulations. Bars show the frequency distributions from 5 nm bins.

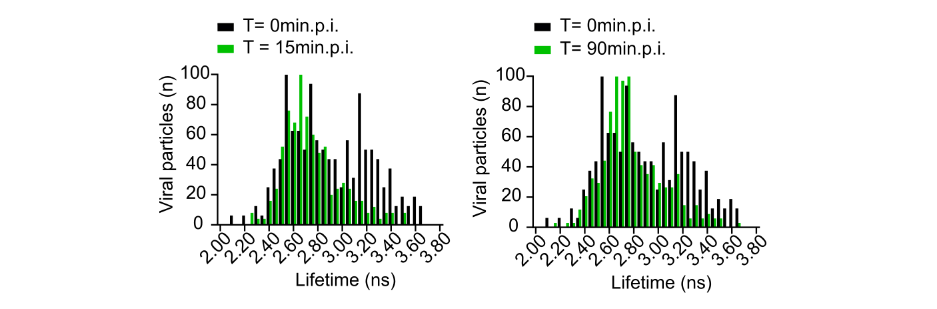

**Supplementary Figure 6. Single-particle fusion assay in Tzm-bl cells.** Frequency distribution of average lifetime values from individual virions containing the Gag-(mScarlet+nirFAST) + HPAR-3,5DOM biosensor. Tzm-bl cells were infected with a 1:1 mixture of viruses produced in absence (mature-enriched) or presence of protease inhibitor (immature, + 500 nM SQV) and time-course of fusion was examined at t=0 (black), 15 min (green, left) and 90 min (green, right) post infection. Bars show frequency of events applying a 0.5 ns bin.
